# Cortical neuroprostheses improve auditory coding and perception compared to cochlear implants

**DOI:** 10.1101/2025.10.27.684780

**Authors:** J.A. Taylor, N. Jovanovic, C.A. Navntoft, A. Albon, S. Rezaei-Mazinani, B. Bathellier, T.R. Barkat

## Abstract

Cochlear implants have transformed the treatment of hearing loss by enabling auditory perception through direct electrical stimulation of the auditory nerve. However, their effectiveness can be limited in noisy environments, for fine frequency resolution, and in patients lacking an intact auditory nerve. The auditory cortex offers a promising alternative target for neuroprosthetic stimulation, but it remains unclear whether direct cortical input can evoke percepts with the complexity and structure of natural sounds. Here we show that surface cortical implants in mice support robust and flexible auditory behaviour, surpassing cochlear implants in tasks requiring fine spectral, temporal, and noise-resistant discrimination. We also show that animals generalize seamlessly between cortical and acoustic stimuli without additional training, indicating that cortical stimulation can generate interpretable, sound-like percepts. Electrophysiological recordings revealed that cortical stimulation drives spatially and temporally structured neural activity in auditory cortex, resembling responses to natural sound. These findings establish that the auditory cortex can interpret spatially and temporally patterned electrical input in a behavioural relevant way. By linking neuroprosthetic stimulation to naturalistic neural representations and behavioural generalization, this work provides a mechanistic and functional foundation for cortical auditory neuroprostheses and points toward new strategies for restoring hearing through direct brain stimulation.

## Main

Hearing loss affects over 430 million individuals globally and is projected to exceed 700 million by 2050^1^. Cochlear implants, the most widely used neuroprosthetic devices, restore partial hearing in patients with severe-to-profound sensorineural loss^2^ (∼1 million patients worldwide with cochlear implants for 50 million cases). Despite their success, cochlear implants suffer from limited spectral resolution and poor sound fidelity, particularly in complex acoustic environments^3,4^. Furthermore, they are ineffective in individuals with damage to the auditory nerve. Although auditory brainstem implants (ABIs) have been developed for such cases, their performance remains modest and inconsistent^5,6^.

The auditory cortex presents a promising alternative site for neuroprosthetic intervention, yet its potential to support meaningful auditory perception remains unclear. Previous work has shown that direct stimulation of auditory cortex can elicit perceptual effects, both in humans and animals. Classic studies by Penfield and colleagues reported that electrical stimulation of auditory cortex during neurosurgery could evoke simple auditory hallucinations, such as tones, buzzing, or voices, providing early evidence for the perceptual relevance of cortical activation^7^. More recently, work in humans demonstrated that stimulation of the planum temporale could enhance speech-in-noise perception^8^, while work in mice showed that optogenetic stimulation of targeted neuronal ensembles in auditory cortex could drive specific neurons and influence decision-making^9^. These studies support the hypothesis that artificial activation of the auditory cortex can generate structured percepts that integrate into meaningful auditory experience.

Here, we assess whether direct electrical stimulation of the auditory cortex can evoke behaviourally relevant auditory percepts in animal models, thereby offering a pathway to expand hearing restoration strategies beyond the cochlea.

### Cortical stimulation yields superior perceptual performance across tasks

To compare auditory perception under cortical and cochlear stimulation with acoustic stimulation, we first established implant models in mice that reflect current clinical and experimental paradigms. For cochlear implantation, we adapted the surgical techniques previously described for acute implantation^10^ and inserted a custom-made four-channel electrode array into the scala tympani of the left cochlea (Fig. 1a, Extended Fig. 1a, b). Functional placement and efficacy were confirmed via auditory brainstem responses (ABRs) recorded pre-implantation and electronic ABRs (eABRs) recorded post-implantation (Fig. 1b).

**Figure 1:**
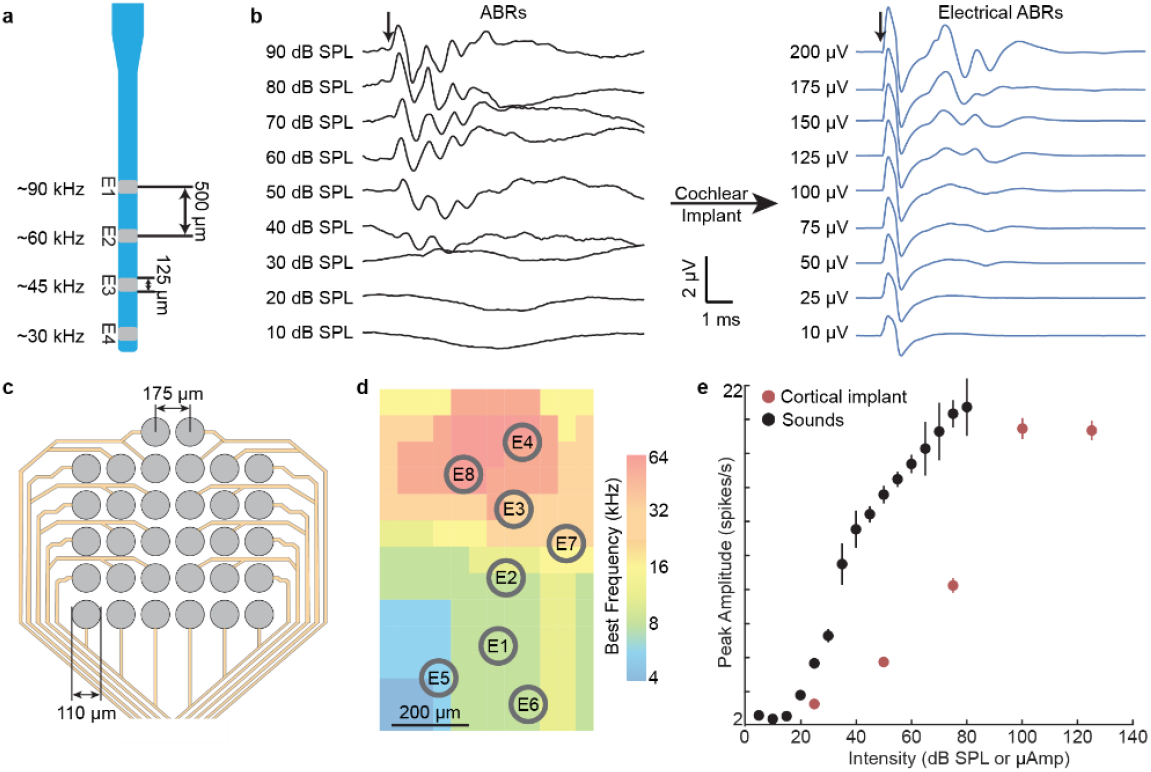
Implant designs and functional validation for auditory stimulation. **a**. Schematic of the four-channel cochlear implant showing 0.500 mm spacing and estimated frequency ranges associated with each electrode. **b**. ABRs to sound clicks before implantation (black traces, left), and eABRs in the same animal following cochlear implantation (blue traces, right). Arrows indicate stimulus onset **c**. Schematic of the 32-channel surface cortical implant with 0.175 mm inter-electrode spacing. **d**. Schematic of tonotopic map of primary auditory cortex (A1) overlaid with cortical implant electrodes, showing alignment with frequency organization. **e**. Peak neuronal response in A1 in response to pure tone sounds (black, n = 5 mice) and electrical cortical stimulation (red, n = 5 mice) as a function of stimulus intensity.

For cortical stimulation, we implanted a previously characterized^11^ 32-channel surface electrode array on top of the primary auditory cortex, positioned above the dura and secured with biocompatible adhesive and dental cement (Fig. 1c, d, Extended Fig. 1c). Prior to implantation, we used penetrating electrodes to map the tonotopic organization of auditory cortex with pure tones, ensuring accurate placement of the array (see methods). The same electrodes were also used to record neural responses to both acoustic stimuli and direct electrical stimulation, confirming functional responsiveness before the surface implant was secured (Fig. 1e). Both cortical and cochlear implants were well tolerated, remained stable over several months, and elicited robust, stimulus-evoked responses in auditory cortex (Extended Fig. 1d, e).

To assess perceptual performance with the implants under different stimulation modalities (acoustic, cochlear or cortical), we trained mice on a series of stimulus discrimination tasks using a go/no-go paradigm (Fig. 2a, Extended Fig. 2). Prior to task onset, animals underwent a detection task, in which they were trained to lick in response to either a single-electrode cortical or cochlear stimulus, or to a single-frequency tone. All animals achieved a ≥75% hit rate in the initial detection task (Extended Fig. 3a–c), ensuring comparable motivation across groups before progressing to the main discrimination tasks. Mice were then trained on the go/no-go paradigm, and performance was tracked daily until a learning plateau was reached, defined as three consecutive days without improvement.

**Figure 2:**
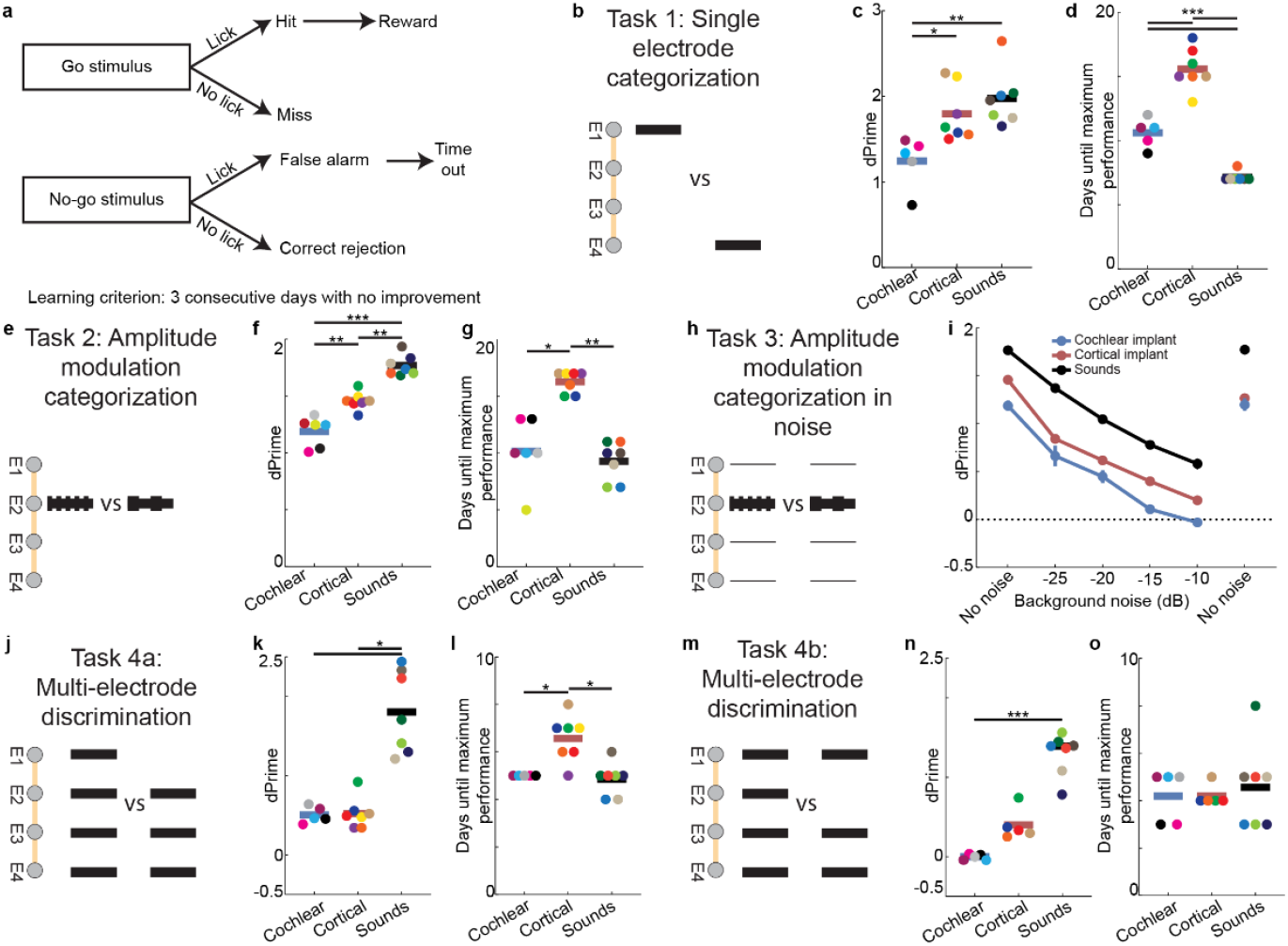
Cortical stimulation yields superior perceptual performance across tasks but slower acquisition than cochlear stimulation. **a**. Schematic of Go/No-go task structure used to assess auditory discrimination in head-fixed mice. **b**. Schematic of task 1: Single-electrode or pure-tone categorization. **c**. Final performance (d′) in task 1 for cochlear, cortical and acoustic stimulations (*, p < 0.05, **, p < 0.01). **d**. Time to reach learning criterion for cochlear, cortical, and acoustic stimulations in task 1 (***, p < 0.001, blue: cochlear implants n = 5 mice, red: cortical implants n = 7 mice, black: sounds n = 7 mice). **e**. Schematic of task 2: Amplitude modulation rate categorization. **f**. Final performance (d′) for cochlear, cortical, and acoustic stimulations in task 2 (**, p < 0.01, ***, p < 0.001). **g**. Time to reach learning criterion for cochlear, cortical, and acoustic stimulations in task 2 (*, p < 0.05, **, p < 0.01, blue: cochlear implants n = 6 mice, red: cortical implants n = 7 mice, black: sounds n = 7 mice). **h**. Schematic of task 3: Amplitude modulation rate categorization in noise. **i**. Performance (d’) in task 3 across different levels of background noise (blue: cochlear implants n = 6 mice, red: cortical implants n = 7 mice, black: sounds n = 7 mice). **j**. Schematic of task 4a: Multi-electrode (frequency pattern) discrimination with edge-omitted (4a). **k**. Final performance (d′) for cochlear, cortical, and acoustic stimulations in task 4a (*, p < 0.05). **l**. Time to reach learning criterion for cochlear, cortical, and acoustic stimulations in task 4a (*, p < 0.05, blue: cochlear implants n = 5 mice, red: cortical implants n = 7 mice, black: sounds n = 7 mice). **m**. Schematic of task 4b: Multi-electrode (frequency pattern) discrimination with centre-omitted (4a). **n**. Final performance (d′) for cochlear, cortical, and acoustic stimulations (***, p < 0.001). **o**. Time to reach learning criterion for cochlear, cortical, and acoustic stimulations in task 4a (*, p < 0.05, blue: cochlear implants n = 5 mice, red: cortical implants n = 5 mice, black: sounds n = 7 mice).

To evaluate frequency discrimination accuracy, the mice learnt to discriminate between individual electrodes or sound frequencies (Task 1, Fig. 2b). Final performance was significantly higher in the cortical group than in the cochlear group (cortical d′ = 1.80 ± 0.12 vs cochlear d’ = 1.24 ± 0.13; p < 0.01; Fig. 2c, Extended Fig. 3d), even though cortical stimulation was delivered via a denser array. The cortical implant featured 175 µm inter-electrode spacing compared to 500 µm for the cochlear array (Extended Fig. 3e), underscoring its capacity to support precise perceptual coding with higher spatial resolution. There was no difference between acoustic and cortical stimulation (acoustic d’ = 1.98 ± 0.12 vs cortical d’ = 1.80 ± 0.12, p >0.99). Interestingly, cochlear-stimulated mice reached criterion faster than cortical-stimulated mice as observed by measuring either the number of days to criterion (11 ± 1 vs. 16 ± 1 days; p < 0.01; Fig. 2d) or the performance on the day cochlea-stimulated mice reached criterion (Extended Fig. 3f). Reaction times did not differ significantly between groups (Extended Fig. 3g).

Amplitude modulations (AM) are an important acoustic feature for sound recognition, particularly speech sounds. To evaluate AM perception with the implants, we trained mice to discriminate between different AM rates (20-200 Hz, see methods) delivered via sound, cochlear stimulation, or cortical stimulation (Task 2, Fig. 2e). Final performance was significantly higher in the cortical than in the cochlear group (d′: 1.46 ± 0.03 vs. 1.19 ± 0.05; p < 0.05, Fig. 2f, Extended Fig. 3h), indicating more precise temporal discrimination. As in Task 1, cochlear-stimulated mice reached criterion faster than cortical-stimulated mice (Fig. 2g, Extended Fig. 3i), again demonstrating faster acquisition but lower endpoint performance. Notably, reaction times were significantly slower in the cochlear group compared to both sound and cortical groups (Extended Fig. 3j), suggesting either increased decisional uncertainty or delayed neural processing under cochlear stimulation.

To evaluate noise resilience, increasing background noise levels were introduced in a stepwise manner across successive sessions of the AM task (Task 3, Fig. 2h). Discrimination performance declined across all groups (Fig. 2i, Extended Fig. 3 k-n). At -15 dB background noise, cochlear-stimulated animals performed at chance (52.0 ± 0.6%), whereas those receiving cortical stimulation or acoustic input maintained above-chance accuracy up to -10 dB noise level (cortical: 52.9 ± 0.5%, acoustic: 60.7 ± 0.6%; p < 0.05; Fig. 2i).

To evaluate the perception of spectral fine structures, we trained mice on a multi-frequency pattern discrimination task (Task 4). Mice distinguished between stimuli containing four frequency components versus stimuli missing either an edge (Task 4a, Fig. 2j) or a middle frequency (Task 4b, Fig. 2m). In Task 4a, performance was comparable across cortical and cochlear groups (d′: 0.50 ± 0.07 vs. 0.40 ± 0.09; n.s.), though both underperformed relative to the sound group (d′: 1.79 ± 0.22; p < 0.05, Fig. 2k). In contrast, for Task 4b, only mice with cortical stimulation achieved above-chance performance (d′: 0.40 ± 0.09 vs. 0.00 ± 0.01 for cochlear; p < 0.05, Fig. 2n), indicating greater sensitivity to internal feature differences in cortical encoding. No significant differences in reaction times were observed across groups (Extended Fig. 3o, p).

Collectively, these findings indicate that while cochlear stimulation supports faster initial learning, cortical stimulation enables more robust and versatile auditory perception, including in conditions where cochlear implants typically show reduced performance, such as noisy environments or acoustically complex sounds.

### Cortical stimulation elicits percepts that are similar to sounds

An open question in the development of cortical auditory neuroprostheses is whether direct electrical stimulation of the auditory cortex can evoke percepts that resemble natural sounds. While our behavioral data demonstrated superior performance with cortical versus cochlear stimulation, it remained unclear whether this advantage reflected task-specific learning or the generation of sound-like percepts. To directly test perceptual equivalence, we assessed bidirectional generalization between cortical stimulation and acoustic input.

Once mice trained exclusively with cortical stimulation reached plateau performance, they were immediately tested on the same tasks using acoustic stimuli, without additional training. Despite the modality switch, animals performed above chance, achieving d′ values of 0.25–0.8 across Tasks 1, 2, and 4 (one sample Wilcoxon test, p < 0.05 for all tasks, chance d’ = 0; Fig. 3; Extended Data Fig. 4a– d). Conversely, mice trained with acoustic stimuli generalized to cortical stimulation, performing above chance across all paradigms with d′ values of 0.62–1.2. Generalization performance decreased modestly relative to the final day of training (average reduction in d′ = 0.72 ± 0.07), likely reflecting minor mismatches in spatial or spectral tuning—for example, current spread or the coarse resolution of cortical tonotopy may produce broader or shifted percepts compared to pure tones. Nevertheless, performance remained robustly above chance, demonstrating that cortical stimulation evokes percepts with sufficient structure and fidelity to support immediate cross-modal transfer.

**Figure 3:**
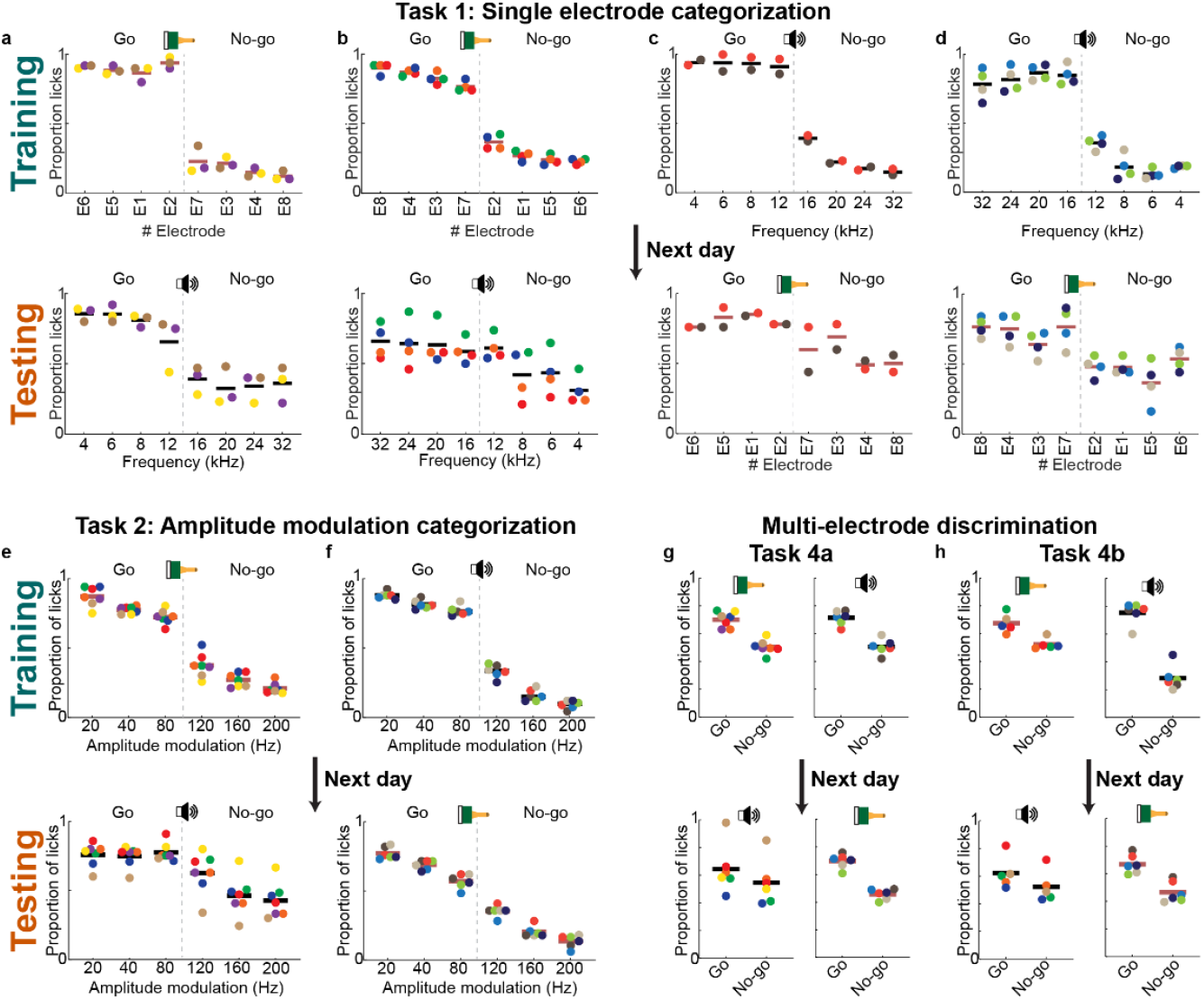
Cortical stimulation elicits percepts that are similar to sounds. **a-d**. Task 1: Final performance of mice trained with cortical stimulation (a, b, top) or sound (c, d, top), and their performance the following day using the opposite modality (bottom) for different batches of mice (a, c: Go stimulation higher frequencies, b, d: Go stimulation lower frequencies). **e, f**. Task 2: Final performance of mice trained with cortical stimulation (e, top) or sound (f, top), and their performance the following day using the opposite modality (bottom). **g**. Task 4a: Final performance of mice trained with cortical stimulation (top left) or sound (top right), and their performance the following day using the opposite modality (bottom). **h**. Task 4b: Final performance of mice trained with cortical stimulation (top left) or sound (top right), and their performance the following day using the opposite modality (bottom).

To confirm that generalization did not arise from rapid relearning, we compared performance during the first and last 100 trials of the test session (Extended Data Fig. 4e–h). Accuracy was stable across this period (First 100 trials average d’ = 0.78 ± 0.08, last 100 trials average d’ = 0.72 ± 0.08) indicating that animals generalized immediately rather than acquiring new associations within the session.

Spectral organization of the percepts was further examined in Task 1 by training separate cohorts with either low or high frequencies/electrodes as the Go stimulus (Fig. 3a, c vs Fig. 3b, d). Performance during training and after modality switching was comparable between groups (final d′ low-frequency Go = 2.1; high-frequency Go = 1.57), indicating consistent perceptual mapping across the spectral axis. A slight reduction in performance for high-frequency Go trials was observed after going from cortical stimulation to sounds, driven primarily by increased false alarms to low-frequency No-go stimuli (false alarms: 44% vs 35%).

Collectively, these results demonstrate that cortical stimulation evokes percepts that are not only discriminable but sufficiently resemble sound to support immediate cross-modal transfer across multiple auditory tasks. This provides behavioural evidence that patterned cortical input can approximate the perceptual structure of hearing, establishing a critical foundation for the development of cortical auditory neuroprostheses.

### Cortical stimulation more faithfully mirrors spatial activation patterns of sound than cochlear stimulation

To examine how cortical and cochlear stimulation engage the auditory cortex at the population level, we performed in vivo electrophysiological recordings in naïve anesthetized mice using a 32-channel silicon probe (four shanks × eight sites) in primary auditory cortex (A1) (Fig. 4a). Recordings were performed in three separate groups receiving either acoustic stimulation, cortical stimulation via surface electrode arrays, or cochlear stimulation via intracochlear electrodes.

**Figure 4:**
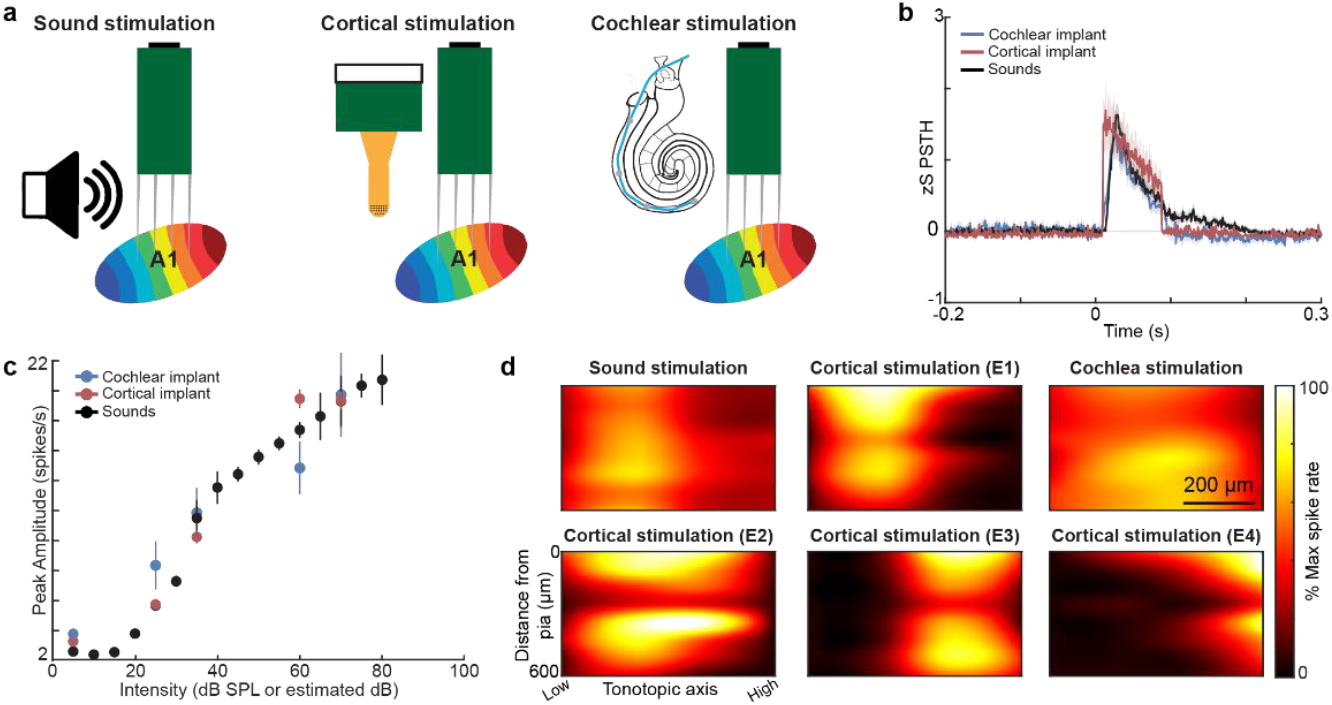
Comparable intensity-dependent responses across modalities but distinct spatial activation profiles in auditory cortex. **a**. Schematic of the *in vivo* electrophysiology setup for recording neural activity across cortical layers using a 32-channel silicon probe in anesthetized mice whilst receiving cochlear, cortical, or acoustic stimulations. **b**. z-scored PSTHs showing average evoked responses across all groups to their respective stimuli. **c**. Average peak response amplitude as a function of stimulus intensity in all conditions (sounds: black, n = 203 from N = 5 mice, cortical: red, n = 149 from N = 5 mice, cochlear: blue, n =129 cells from N = 5 mice). **d**. Example heatmaps of cortical activity across electrode depth and shank position in response to a single electrode cochlea stimulation (top right), individual electrode cortical surface stimulation (top centre and bottom), and pure tone sound stimulation (top left).

Across all groups, we observed robust evoked activity in response to their respective stimuli, as shown in population peristimulus time histograms (PSTHs; Fig. 4b). Peak response amplitude increased with stimulus intensity in all conditions (Fig. 4c, see methods for estimation of dB), consistent with scalable encoding of input strength in auditory cortex^12^.

To examine spatial coding, we analysed depth-resolved activity in response to single pure tones or single-electrode stimulation. Acoustic stimulation evoked a canonical columnar activation pattern centred in deeper cortical layers (Fig. 4d, top left, Extended Fig. 5a), reflecting thalamocortical input^13^. Cortical stimulation produced a similar columnar profile with additional strong activation at the pial surface (Fig. 4d, top centre and bottom, Extended Fig. 5b), consistent with current injection from surface electrodes. Cochlear stimulation also generated columnar activation but exhibited broader lateral spread (Fig. 4d, top right, Extended Fig. 5c), potentially reflecting current spread in the cochlea and contributing to the poorer discrimination seen in tasks 1 and 4. Notably, spatial resolution under cortical stimulation was sufficiently fine that distinct electrodes elicited topographically shifted response columns along the tonotopic axis of auditory cortex (example in Fig. 4d, bottom row, Extended Fig. 5b). This spatial specificity likely supports the superior perceptual performance observed under cortical stimulation, especially for tasks requiring fine spectral discrimination.

### Cortical stimulation recapitulates sound representations in neuronal population space better than cochlear implants

To determine how different stimulation modalities are represented in auditory cortex, we examined neuronal activity across all behavioural paradigms using a combination of supervised decoding, representational similarity analysis, and dimensionality reduction (Fig. 5). For each task, a two-way support vector machine (SVM) was trained to distinguish spontaneous from evoked activity patterns in single units, providing a common measure of how effectively each modality drove distinct temporal patterns in the auditory cortex. Decoders trained on sound-evoked responses generalized to cortical stimulation with high accuracy across all tasks (Proportion correct = 77%, 72%, 67% tasks 1, 2 and 4 respectively, Fig 5a, e, h, k, n), whereas generalization to cochlear stimulation was consistently lower (Proportion correct = 60%, 69%, 61%, tasks 1, 2 and 4 respectively, Fig 5a, e, h, k, n). The same pattern emerged when training on cortical stimulation, with sound responses matching within-modality accuracy but cochlear responses performing worse (Extended Fig. 6e, proportion correct sound: 69%, cortical: 70%, cochlear: 63%) In contrast, decoders trained on cochlear responses generalized poorly to both other modalities (Extended Fig. 6f, proportion correct sound: 54%, cortical: 50%, cochlear: 66%).

**Figure 5:**
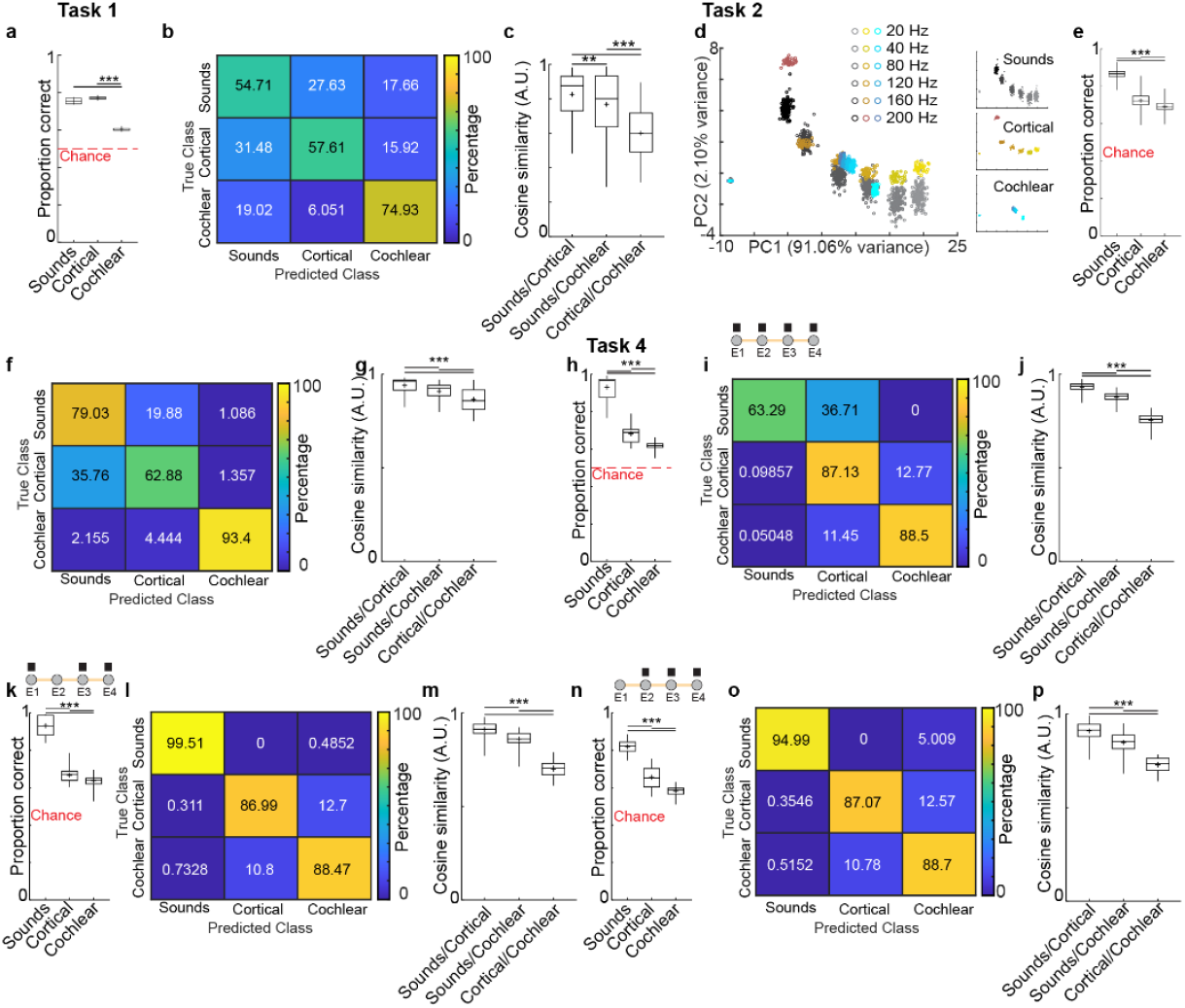
Cortical stimulation elicits population-level neural responses in auditory cortex that more closely resemble sound-evoked patterns than cochlear stimulation. **a**. Classification accuracy of a two-way SVM trained to distinguish spontaneous from sound-evoked activity (n = 142 cells), and tested on held-out sound responses (n = 60 cells), cortical stimulation responses (n = 149 cells), and cochlear stimulation responses (n = 129 cells, bootstrap, p < 0.01) **b**. Confusion matrix of a three-way SVM trained to classify neuronal responses as originating from acoustic, cortical, or cochlear stimulation (training: n = 337 cells, testing n = 144 cells, bootstrap, p < 0.01). **c**. Cosine similarity between trial-averaged auditory cortical population responses evoked by acoustic, cortical or cochlear individual electrode/frequency stimulation (Kruskal-Wallis multiple comparisons test) **d**. PCA of responses to amplitude-modulated (AM) stimuli delivered via sound (black, n = 203 cells from N = 5 mice), cortical stimulation (red, n = 149 cells from N = 5 mice), or cochlear stimulation (blue, n = 129 cells from N = 5 mice). **e**. Classification accuracy of a two-way SVM decoding accuracy trained to distinguish spontaneous from sound-evoked AM activity (n = 142 cells), and tested on held-out sound responses (n = 60 cells), cortical stimulation responses (n = 149 cells), and cochlear stimulation responses (n = 129 cells, bootstrap, p < 0.01). **f**. Confusion matrix from three-way SVM classification of AM responses (training: n = 337 cells, testing n = 144 cells, bootstrap, p < 0.01). **g**. Cosine similarity between trial-averaged auditory cortical population responses evoked by acoustic, cortical or cochlear AM stimulation (Kruskal-Wallis multiple comparisons test). (**h, k, n**) Classification accuracy of a two-way SVM decoding accuracy trained to distinguish spontaneous from sound-evoked multi-electrode activity (n = 142 cells), and tested on held-out sound responses (n = 60 cells), cortical stimulation responses (n = 149 cells), and cochlear stimulation responses (n = 129 cells, bootstrap, p < 0.01). (**i, l, o**). Confusion matrix from three-way SVM classification of multi-frequencies or multi-electrode stimuli for four electrodes (i), centre-omitted (l) and edge-omitted (o) (training: n = 337 cells, testing n = 144 cells, bootstrap, p < 0.01). (**j, m**, **p**) Cosine similarity between trial-averaged auditory cortical population responses evoked by acoustic, cortical or cochlear multi-electrode/frequency stimulation (Kruskal-Wallis multiple comparisons test).

Three-way classifiers were then trained to discern between single unit responses to the different modalities. The results further confirmed this representational hierarchy, more often confusing cortical and sound responses than either was with cochlear responses (bootstrapped p < 0.01, Fig. 5b, f, i, l, o and Extended Fig. 6h, 7d, 8b, 9b). Representational similarity analysis revealed the same trend: cosine similarity was highest between cortical and sound responses (Task 1: 0.82, Task 2: 0.91, Task 3: 0.79, Task 4: 0.92, Fig. 5c, g, j, m, p and Extended Fig. 8c), intermediate for sound–cochlear (Task 1: 0.76, Task 2: 0.88, Task 3: 0.64, Task 4: 0.86, Fig. 5c, g, j, m, p and Extended Fig. 8c), and lowest for cortical– cochlear (Task 1: 0.60, Task 2: 0.86, Task 3: 0.65, Task 4: 0.73, Fig. 5c, g, j, m, p and Extended Fig. 8c). PCA and T-SNE visualizations (Fig. 5d, Extended Fig. 6b-d, 7b, 8a, 9a) provided a complementary perspective, showing closer clustering of cortical and sound responses across all tasks, from frequency discrimination to complex pattern recognition.

This convergence between cortical and acoustic representations was robust across paradigms with differing sensory demands, whether resolving fine frequency steps, discriminating amplitude modulation rates, or identifying patterns in noise, and it paralleled the behavioural outcomes. Given that mice trained with cortical stimulation showed superior resilience to background noise in the amplitude modulation task, the next step was to ask whether this behavioural advantage is reflected in population-level neural coding. We therefore examined whether cortical stimulation, like natural sound, supports precise temporal encoding of modulation rates and whether this fidelity persists under increasing noise, using a neurometric analogue of the behavioural paradigm.

### Neuronal decoding mirrors behavioural performance

To directly link neural population coding with perceptual performance, we trained six-way SVM classifiers on auditory cortical responses to the amplitude modulation (AM) stimuli used in Tasks 2 and 3 (Fig. 6). Classifiers were tasked with identifying one of six AM rates based on single-trial neuronal population activity, with decoding accuracy serving as a proxy for neural discriminability under each stimulation modality and noise condition.

**Figure 6:**
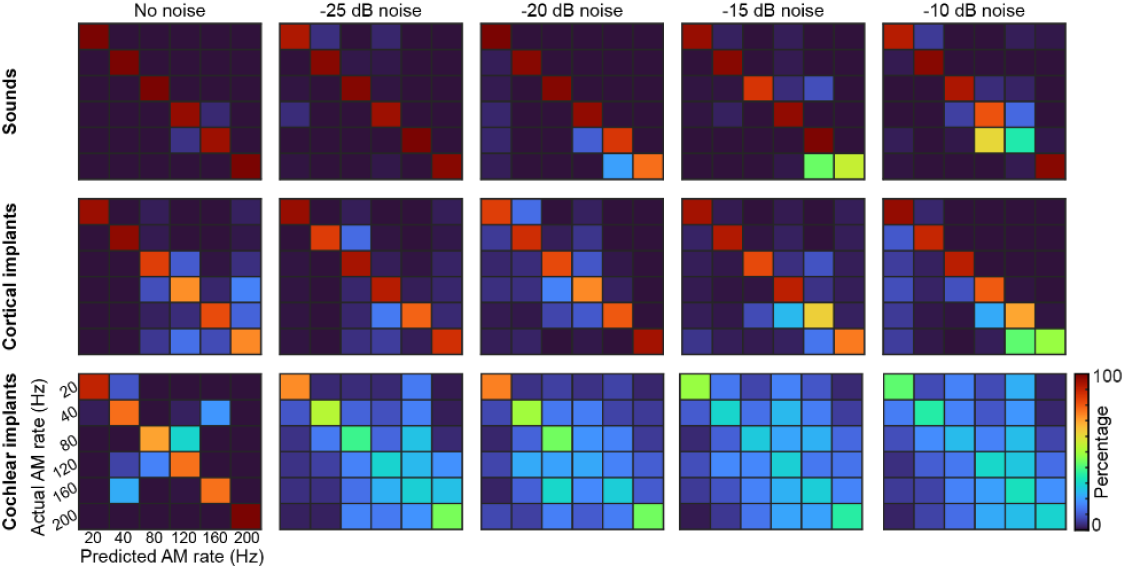
Neural decoding of amplitude modulation rate mirrors behavioural performance. Six-way SVM decoders were tested for each group to discriminate between different rates of amplitude modulation under different noise conditions (Sounds: training n= 142 cells, testing n = 61 cells. Cortical implants: training n = 104 cells, testing n = 45 cells. Cochlear implants: training n = 90 cells, testing n = 39 cells).

In the absence of background noise, decoder performance for acoustic stimuli was near perfect, with only minor confusion between adjacent high modulation rates (e.g., 3.5% error between 120 and 160 Hz). Accuracy remained high at -10 dB SPL background noise (83%), indicating the resilience of sound-evoked cortical representations. Cortical stimulation also yielded high classification accuracy in quiet (85%) and remained robust under increasing noise, with only a modest decline to 79% at -10 dB SPL noise. These results closely paralleled behavioural performance for both modalities.

In contrast, cochlear stimulation produced less reliable population-level coding of modulation rate. While baseline accuracy in quiet conditions was relatively high (82%), performance deteriorated rapidly with added noise. At -25 dB SPL background noise, accuracy dropped to 45%, approaching chance level (16.7%), and further declined to 32% at -10 dB SPL. This pattern closely mirrors behavioural data, where cochlear-implanted animals failed to perform the AM-in-noise task above -20 dB SPL background.

Together, these findings demonstrate that cortical and acoustic stimulation engage population-level coding schemes capable of supporting precise temporal discrimination, even under noisy conditions. The alignment between neurometric and behavioural performance underscores the functional viability of cortical stimulation as a robust and perceptually meaningful auditory interface.

## Discussion

Cochlear implants have transformed hearing restoration, but limitations in spectral resolution and reliance on the auditory nerve leave some patients without effective options. Our findings demonstrate that direct stimulation of the auditory cortex via surface implants can enable robust auditory perception in mice. Across multiple tasks, ranging from frequency discrimination and temporal patterning to noise resilience, cortical stimulation outperformed cochlear stimulation. Notably, animals trained with cortical implants generalized bidirectionally to and from sound-based tasks without additional training, providing strong evidence that cortical stimulation evokes percepts with sound-like structure.

These behavioural outcomes were mirrored at the neuronal level. Electrophysiological recordings revealed that cortical stimulation elicited auditory cortical responses that closely resembled those evoked by natural sounds in terms of spatial activation patterns, dimensionality, and decodability. Compared to cochlear stimulation, cortical input preserved temporal and spectral features of sound more faithfully, especially under challenging conditions like background noise. These results position cortical neuroprostheses as a viable alternative for hearing restoration, especially for individuals with damage to the auditory nerve.

The enhanced performance of cortical stimulation in frequency discrimination may be rooted in the intrinsic properties of auditory cortical processing. In the human auditory cortex, single neurons exhibit frequency tuning that is markedly narrower than that observed in the periphery, indicating that spectral resolution in cortex exceeds that available from the auditory nerve^14^. This hyperacuity is likely shaped by nonlinear mechanisms that go beyond simple frequency selectivity and contribute to fine-grained stimulus encoding. Consistently, bilateral cortical lesions lead to elevated frequency discrimination thresholds that approximate peripheral tuning, suggesting that normal cortical function is required to extract the full precision of frequency information represented subcortically^15^. In this context, the superior frequency discrimination observed with cortical implants may reflect the ability of spatially patterned stimulation to tap into this cortical-level refinement, while cochlear implants remain constrained by the broader tuning and coarser spatial representation of the auditory nerve.

The learning dynamics observed between the two modalities offers important insights. Mice using cortical stimulation took significantly longer to reach peak performance compared to those with cochlear implants. One possible explanation is the lack of upstream subcortical input during cortical stimulation, which may limit the natural feedforward-driven plasticity in auditory circuits. Another contributing factor may be how stimulation strength was calibrated: cochlear implants were matched to auditory brainstem responses, while cortical implants were matched directly to A1 activity. Because coding principles differ between the cochlea and cortex, particularly in how intensity and temporal features are represented, this calibration is not perfectly equivalent, which could also influence learning trajectories and final performance. Approaches to accelerate learning could include neuromodulation-based plasticity enhancement, shown to be effective in cochlear implant models^16^, or even dual implant strategies that combine cochlear and cortical stimulation to leverage the strengths of both systems.

One limitation of this study is its focus on a mouse model. Nevertheless, our results resonate with previous human studies. Electrical stimulation of the auditory cortex, particularly Heschl’s gyrus, has been shown to elicit auditory percepts^17^. Additionally, the topography and content of these percepts have been found to align with natural sound-evoked patterns, suggesting that the human cortex is capable of interpreting spatially patterned input as meaningful auditory information^18^. These observations raise the exciting possibility that high-density cortical arrays could support discrete, intelligible auditory percepts in clinical settings.

Our findings also align with broader efforts to replace missing sensory input through direct brain stimulation. In vision restoration, intracortical stimulation of the visual cortex using Utah arrays has enabled blind patients to perceive borders, shapes, and even characters^19^. In parallel, innovations in ABIs using flexible high-density micro-electrodes similar to the ones used in this study have shown the potential to improve perceptual outcomes^20^, further underscoring the importance of spatial precision in neuroprosthetic design. Here, we deliver patterned input through a high-density surface array with contacts less than 200 µm apart, enabling fine-grained spatial activation without penetrating the cortex. This approach offers a unique combination of resolution, coverage, and minimal invasiveness and represents a substantial advance over previous human cortical stimulation efforts in which surface electrodes were spaced on the millimetre scale^21^.

While electrical stimulation at various levels of the auditory pathway has been explored, including at the cochlea and brainstem, existing approaches have either depended on intact peripheral structures or failed to evoke perceptually meaningful representations. Our findings provide a convincing demonstration that surface cortical stimulation alone can drive robust auditory behaviour, bidirectional generalization with natural sound, and neural population activity that recapitulates acoustic coding structure. This establishes that the auditory cortex can interpret spatially patterned electrical input in a perceptually relevant way, even without prior engaging subcortical processing. By showing that naturalistic and transferable auditory percepts can be evoked through nonpenetrating cortical interfaces, this work marks a significant step toward next-generation neuroprosthetic solutions.

## Methods

### Animal use and ethics

All animal procedures were conducted in accordance with the University of Basel’s institutional guidelines and approved by the Veterinary Office of the Canton Basel-Stadt, Switzerland. Experiments were carried out on male and female C57Bl/6JRj mice (Janvier Labs, France), aged 6–8 weeks at the time of surgery. Mice were housed under a 12-hour light/dark cycle with *ad libitum* access to food and water. Following surgery, animals remained single-housed with unrestricted access to food. During behavioral training, food intake was manually regulated starting the day before training onset, and body weight was monitored daily to maintain animals between 85% and 95% of baseline. All experiments were performed during the light phase.

### Surgical protocols

Mice were anesthetized with ketamine (80 mg/kg) and xylazine (16 mg/kg), administered intraperitoneally, and received local analgesia via a subcutaneous injection of bupivacaine (0.01 mg/animal) and lidocaine (0.04 mg/animal). Supplemental ketamine (45 mg/kg) was provided as needed during surgery. Perioperative analgesia was administered via intraperitoneal buprenorphine (0.1 mg/kg). Body temperature was maintained at 37 °C using a heating pad (FHC, ME, USA), and ophthalmic lubricant was applied to both eyes.

#### Chronic cochlear implants

ABRs were recorded by occluding the contralateral ear with acoustic foam and inserting subdermal electrodes at the vertex (active), below the ipsilateral pinna (reference), and in the hindlimb (ground). Click stimuli were generated using a digital signal processor (RZ6, Tucker Davis Technologies) at a 200 kHz sampling rate and delivered through a magnetic speaker (MF1, Tucker Davis Technologies, FL, USA) positioned 10 cm from the left ear. Stimuli were presented in 10 dB SPL increments from 90 to 10 dB SPL and averaged across 512 repetitions per level. Recordings were band-pass filtered (300– 2,000 Hz), and auditory thresholds were determined based on response amplitude.

Following ABR recordings, a four-channel cochlear implant (Oticon medical) was inserted as previously described^10^. A subcutaneous ground wire was positioned in the neck, and the implant connector was affixed to the skull with Loctite adhesive and dental cement, along with a custom headpost. The implant wire is 200 µm diameter and 14 mm long with four ring electrodes of 125 µm separated by 500 µm center to center.

eABRs were recorded using the same electrode configuration. Single bipolar pulses (10 ms) were delivered via a stimulus isolator (IZ2, Tucker Davis Technologies, FL, USA) at amplitudes ranging from 10 to 200 μV in 5 μV increments. Recordings were band-pass filtered (300–2,000 Hz), and auditory thresholds were determined based on response amplitude. These recordings enabled alignment of electrical and acoustic input intensities by comparing ABR and eABR thresholds.

#### Chronic cortical implants

A craniotomy was performed over the auditory cortex, and the exposed brain surface was covered with silicone oil to prevent drying. A multi-channel silicon probe (A4x8-5mm-50-200-177-A32, NeuroNexus) was inserted using a motorized stereotaxic micromanipulator (DMA-1511, Narishige, Japan) to functionally localize A1 based on its tonotopic organization, characterized by a caudo-rostral increase in best frequency. Following A1 identification, the probe was withdrawn and a custom-designed 32-channel cortical electrode array^11^ was placed on the cortical surface above the dura, aligned to the functionally mapped A1 region. The cortical implant consisted of a 32-channel electrode array, with circular electrodes 110 µm in diameter spaced 175 µm center-to-center. Electrodes were fabricated from the conductive polymer PEDOT:PSS to reduce impedance and improve charge transfer. Conductive traces were made of gold, arranged in two stacked layers separated by parylene-C as the insulating substrate. This multilayer configuration enabled high-density wiring while maintaining flexibility and durability of the device.

To secure the implant and protect the underlying brain tissue, the craniotomy was sealed with a silicone elastomer (Kwik-Sil, World Precision Instruments, FL, USA). The array connector and a custom-fabricated headpost were affixed to the skull using Loctite adhesive and dental cement.

A1 neuronal responses to cortical stimulation were recorded at 25 µv intervals of intensity. These recordings enabled alignment of electrical and acoustic input intensities by comparing A1 responses to acoustic and cortical electrical stimuli.

### Behavioural training

Following surgery, mice were allowed to recover for a minimum of 10 days and until they regained their pre-operative weight and showed no signs of post-operative stress. Food restriction began 24 hours prior to the start of training. During the first day of habituation, mice were head-fixed for one hour with free access to the lick port for food reward. On the second day, they began the detection task.

Auditory stimuli were generated using a digital signal processor (RZ6, Tucker Davis Technologies, FL, USA) at a 200 kHz sampling rate and delivered through a calibrated MF1 speaker (Tucker Davis Technologies) positioned 10 cm from the left ear. Calibration was performed with a wide-band ultrasonic acoustic sensor (Model 378C01, PCB Piezotronics, NY, USA), and sound levels were set to 70 dB SPL. Electrical stimuli were generated and delivered through the same processor and a stimulus isolator (IZ2, Tucker Davis Technologies), calibrated to evoke responses equivalent to 70 dB SPL based on ABRs or cortical recordings in A1. For cochlear implants, stimulation levels were calibrated by matching ABRs evoked by acoustic clicks to eABRs obtained at varying current levels of cochlear stimulation. For cortical implants, calibration was performed by aligning sound-evoked responses in primary auditory cortex with those elicited by electrical cortical stimulation, ensuring comparable response amplitudes across modalities.

In the initial detection task, mice learned to associate a single stimulus (sound or electrical) with a reward. Once they achieved ≥75% hit rates, they progressed to the Go/No-go discrimination tasks. Each behavioral session consisted of 400 trials (200 Go, 200 No-go) and lasted approximately one hour, with one session per day.

Task 1 (Frequency/Electrode Discrimination): Mice first learned to discriminate between two frequencies (sound group) or two electrodes (implant groups). Upon reaching ≥75% correct responses, task complexity increased to four cochlear electrodes or eight cortical electrodes/frequencies. For between-group comparisons, only four linearly spaced stimuli were analyzed. Mice trained until reaching a behavioral plateau, defined as three consecutive days with < 0.2 change in d′.

Task 2 (Amplitude Modulation Discrimination): Mice initially discriminated between 20 Hz and 200 Hz modulated stimuli. Once ≥75% accuracy was reached, they were presented with six AM rates: 20, 40, and 80 Hz (Go), and 120, 160, and 200 Hz (No-go). Training continued until d′ performance plateaued for three consecutive days.

Task 3 (AM in Noise): The same AM task as Task 2 was performed with added background noise. For the implant groups, noise was introduced by stimulating three additional electrodes at a constant intensity; for the sound group, white noise was played concurrently. One session was completed per noise level which increased each day until -10 dB noise level. To rule out motivation-related effects, animals were tested again without noise the following day. For cochlear implants the noise levels were estimated with the alignment of ABRs and eABRS. For cortical implants the noise levels were estimated with the alignment of response in A1 to acoustic and cortical electrical stimulation.

Task 4 (Multi-Frequency Pattern Discrimination): Mice learned to distinguish between full patterns (four electrodes/frequencies) and stimuli with one component missing (edge or center). Training continued until a behavioral plateau was reached.

Generalization across stimulation modalities: To evaluate perceptual generalization between stimulation modalities, mice trained with either cortical stimulation or sound were tested using the alternate modality at the end of each task. Specifically, after completing Task 1 with their trained modality, animals were presented with the same task using the other modality (e.g., sound-trained mice received cortical stimulation, and vice versa) before progressing to Task 2. This cross-modal testing approach was repeated for Tasks 2 and 4, enabling direct assessment of perceptual equivalence without additional training between modality switches.

### Electrophysiology

Extracellular recordings were performed in anesthetized, head-fixed, naïve mice inside a sound-attenuating chamber (MAC-2, Industrial Acoustics Company Nordics, Hvidovre, Denmark). For the cochlear implant group, the location of A1 was functionally mapped prior to cochlear implantation. After insertion of the implant, a multi-channel silicon probe was lowered into A1 for recordings. For the cortical implant group, A1 was similarly mapped before positioning the surface cortical implant; the recording probe remained in place during stimulation. For the sound group, A1 mapping was followed directly by sound stimulation and recording. In all cases, auditory stimuli matched those used in behavioral tasks and were presented in the same order. At the conclusion of recordings, mice were euthanized via intraperitoneal injection of pentobarbital followed by cervical dislocation.

### Data analysis

#### Behavioral performance analysis

Discrimination performance was quantified using proportion of licks to individual stimuli (Fig. 3, extended Fig. 3) and/or *d′* (d-prime, Fig. 2, extended Fig. 4), a measure from signal detection theory that reflects an animal’s sensitivity to distinguish between Go and No-Go stimuli. *d′* was calculated as:

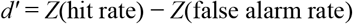

where *Z* is the inverse cumulative distribution function of the standard normal distribution. The hit rate was defined as the proportion of correct responses to Go stimuli, and the false alarm rate as the proportion of incorrect responses to No-go stimuli.

#### In vivo electrophysiology analysis

Extracellular signals were acquired using a 32-channel recording system (RZ2 Bioamp Processor, Tucker Davis Technologies) at 24,414 kHz. Spike sorting was performed offline with KiloSort 2.5 (CortexLab, UCL), followed by manual curation using Phy based on waveform shape, signal-to-noise ratio, and auto-/cross-correlograms. To eliminate stimulation artifacts, a 2 ms blanking window was applied around each individual pulse, and any residual artifacts were removed through manual curation. Both single and multi-units were included in downstream analyses. Further processing was conducted using custom MATLAB R2020b (MathWorks, MA, USA) scripts.

#### Spatial responses analysis

To visualize the spatial distribution of stimulus-evoked activity in A1 (Fig. 4d), we analysed spike responses across the 32-channel silicon probe (four shanks × eight sites). For each stimulus, PSTHs were computed from −0.2 s to +1.2 s relative to stimulus onset, with the 1 s stimulus duration centred in the window. Spike counts were z-scored relative to the baseline period (−0.2 to 0 s) to normalize across units and highlight stimulus-driven changes. To generate continuous spatial activity maps, responses were interpolated using MATLAB’s interp2 function. No averaging across animals was performed; instead, representative examples were selected to illustrate laminar and columnar response patterns under each stimulation modality.

#### Dimensionality reduction analysis

To examine the structure of neuronal population responses across stimulation modalities, we applied dimensionality reduction techniques to trial-averaged firing patterns. For each unit, PSTHs were computed from −0.2 s before stimulus onset to +0.2 s after stimulus offset, encompassing the full 1 s stimulus duration. PSTHs were binned at 10 ms resolution and z-scored to normalize activity across units. The resulting vectors were used as input to PCA and t-distributed stochastic neighbour embedding (t-SNE) for visualization. Dimensionality reduction was performed separately for each behavioural task and modality. PCA was used to assess variance structure, while t-SNE was used to qualitatively visualize clustering in low-dimensional space.

#### Unsupervised learning analysis

To assess the similarity and discriminability of neuronal population responses across modalities, we trained SVM classifiers on trial-averaged activity using custom MATLAB code. Spike trains were binned in 10 ms intervals and z-scored per unit. All classification analyses were repeated over 1,000 random subsamplings to ensure robustness.

Two-way decoders were trained to distinguish between 1 s epochs of spontaneous and stimulus-evoked activity of single units. The feature space was the trial-averaged activity of the unit sampled in 100 bins of 10 ms. The SVM was trained on 70% of units recorded under one modality (e.g., sound), and tested on the held-out 30% of that group, as well as on units from other modalities (e.g., cortical or cochlear stimulation). Accuracy on out-of-modality data was used to quantify cross-modal generalization.

Three-way decoders were trained to classify responses by modality (sound, cortical, cochlear) using single unit activity sampled in 10 ms bins from −0.2 s before to +1.2 s after stimulus onset. The classifier was trained on 70% of all units and tested on the remaining 30%. A confusion matrix was computed to assess patterns of misclassification across modalities.

Six-way decoders were used for amplitude modulation (AM) rate classification. SVMs were trained separately for each modality to distinguish among six AM rates (20, 40, 80, 120, 160, 200 Hz), using single unit activity sampled into 10 ms bins from −0.2 s to +1.2 s relative to stimulus onset. Performance was measured as mean classification accuracy across iterations.

All classifiers used linear kernels and default MATLAB hyperparameters unless otherwise specified.

### Statistics

All statistical analyses were performed using MATLAB R2020b (MathWorks, MA, USA) or GraphPad Prism 9 and 10 (GraphPad Software, MA, USA). No statistical methods were used to predetermine sample sizes, but group sizes were consistent with prior studies in the field. Comparisons across three or more groups were conducted using the non-parametric Kruskal–Wallis test, followed by Dunn’s post hoc multiple comparisons. Decoder performance was evaluated using non-parametric bootstrap resampling to estimate confidence intervals and statistical significance. For each condition, decoder accuracy was calculated across 1000 independent iterations of randomized train–test splits. To estimate the uncertainty around mean accuracy, we performed bootstrap resampling with replacement (n = 1000), drawing with replacement from the set of accuracy values obtained across iterations. The resulting bootstrap distributions represent the sampling variability of the mean decoder performance. Two-sided p-values were computed by quantifying the proportion of bootstrap samples with mean accuracy less than or equal to chance level (e.g., 33.3% for three-way classification or 50% for binary classification). Significance was defined at p < 0.05 (*), p < 0.01 (**), and p < 0.001 (***), corresponding to 95%, 99%, and 99.9% confidence intervals, respectively.

Data collection and analysis were not performed blind to experimental condition; however, all analyses were conducted using standardized, automated code applied uniformly across datasets. All animals were included unless they failed to meet learning criteria—defined as fewer than 75% correct responses on a task after 10 training sessions.

## Acknowledgements

This work was funded by the European Union’s Horizon 2020 research and innovation program under grant agreement No 964568 (Hearlight) (to S.R-M., B.B. and T.R.B.). J.A.T. was funded by the Forschungsfonds Nachwuchsforschende of the University of Basel.

## Author information

Authors and affiliations

### Department of Biomedicine, University of Basel, Switzerland

James Alexander Taylor, Natasa Jovanovic, Charlotte Amalie Navntoft & Tania Rinaldi Barkat

### Department of Bioelectronics, Ecole des Mines de Saint-Etienne, France

Amelie Albon & Shahab Rezaei-Mazinani

### Institut de l’audition, Institut Pasteur, Paris, France

Brice Bathellier

### Contributions

J.A.T and T.R.B designed the experiments. J.A.T and N.J. performed the experiments and acquired data. J.A.T. analyzed the data. J.A.T., C.N., A.A. and S.R-M. designed, developed and/or fabricated hardware. J.A.T, B.B. and T.R.B. designed the study. J.A.T and T.R.B wrote the paper.

