## Extended data for "Cortical neuroprostheses improve auditory coding and perception compared to cochlear implants"

### Extended data figures

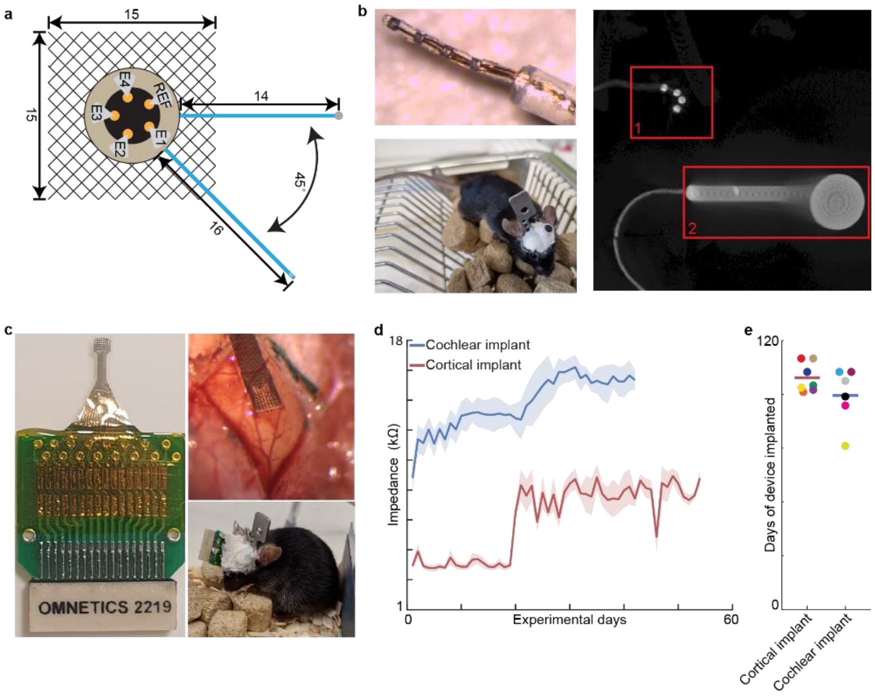


**Extended figure 1: Chronic implants in mice**

a. Schematic of chronic cochlear implant. b. (Top left) Photo of the four electrodes from the cochlear implant. (Bottom left) Photo of mouse with chronic cochlear implant. (Right) X-ray of chronic cochlear implant in mouse. Box 1 shows the four electrodes curving in the cochlear. Box 2 shows the connector of the implant. c. (Left) Photo of chronic cortical implant. (Top right) Photo of cortical implant on A1). (Bottom right) Photo of mouse with chronic cortical implant. d. Impedance of electrodes across experimental days (blue, cochlear implant n = 24 electrodes from N = 6 devices. Red, cortical implants n = 224 electrodes from N = 7 devices). e. Number of days the devices were implanted (Red, Cortical implants N = 7 devices. Blue, Cochlear implants N = 6 days).

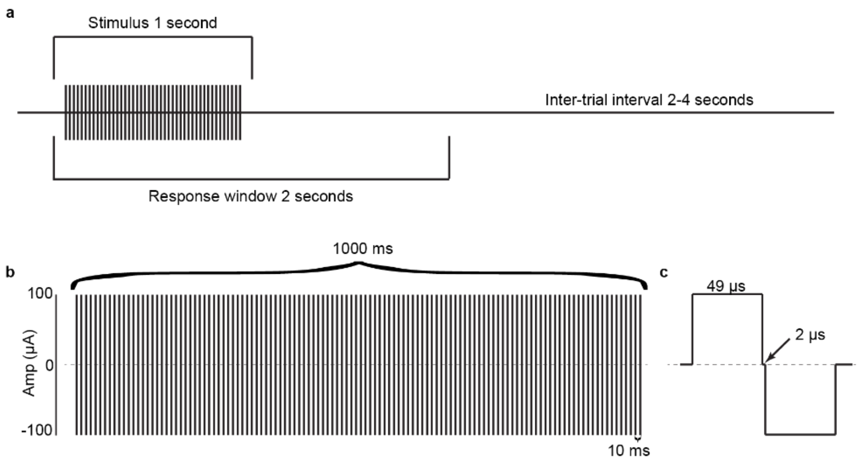


**Extended figure 2: Structure of Go/No-go paradigm and electrical stimulation parameters**

a. Diagram of an individual trial of Go/No-go paradigm. b. Diagram of the electrical stimulation protocol used for both cochlear and cortical implants: each stimulus consisted of 100 biphasic pulses at 100 Hz of 1 ms. c. Diagram of an individual biphasic pulse showing charge-balanced stimulation.

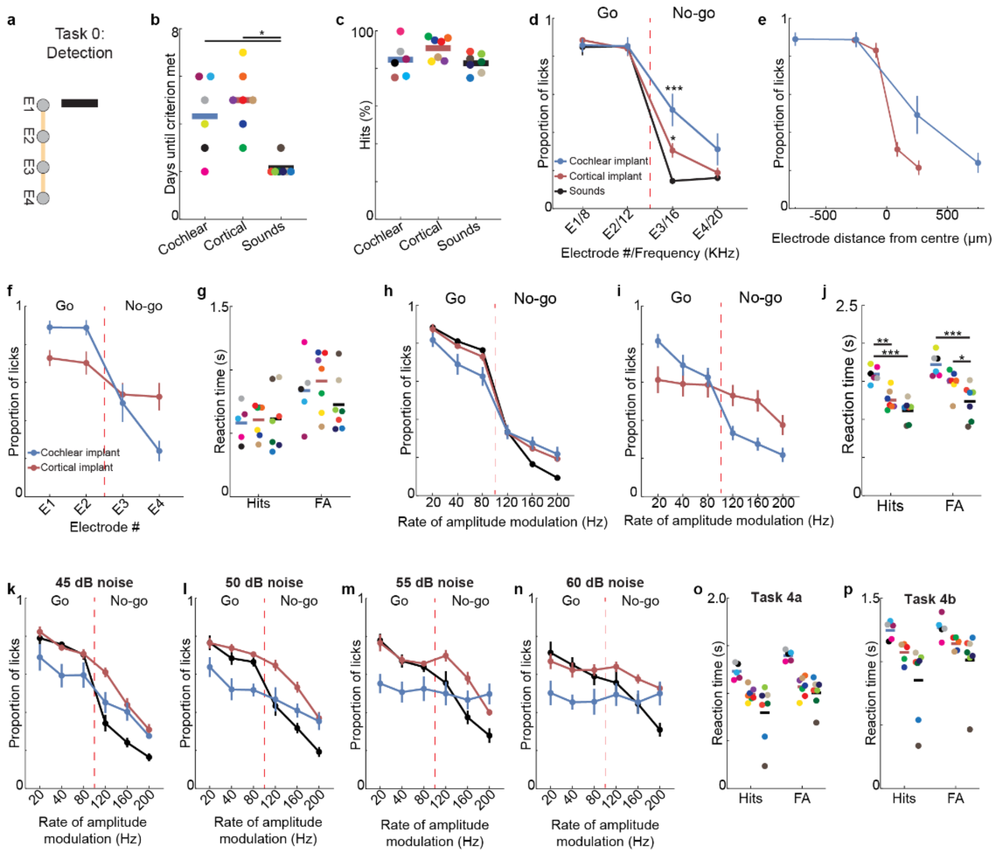

**Extended figure 3: Detection performance, learning dynamics and reaction times**

a. Schematic of detection paradigm. b. Time to reach learning criterion (>75% hits) for acoustic, cochlear, and cortical stimulations (*, $p < 0.05$). c. Performance of mice to detection task (blue: cochlear implants n = 5 mice, red: cortical implants n = 7 mice, black: sounds n = 7 mice). d. Proportion of licks to individual electrodes/ frequencies in task 1 (blue: cochlear implants n = 5 mice, red: cortical implants n = 7 mice, black: sounds n = 7 mice). e. Proportion of licks for individual electrodes based on electrode distance from the centre of the implant. f. Proportion of licks for individual electrodes on day 11, when the mice reached their maximum performance with the cochlear implants. g. Reaction times to hits and FA in task 1 (blue: cochlear implants n = 5 mice, red: cortical implants n = 7 mice, black: sounds n = 7 mice). h. Proportion of licks to different rates of amplitude modulation (blue: cochlear implants n = 6 mice, red: cortical implants n = 7 mice, black: sounds n = 7 mice). i. Proportion of licks to different rates of amplitude modulation on day 10, when the mice reached their maximum performance with the cochlear implants (blue: cochlear implants n = 6 mice, red: cortical implants n = 7 mice, black: sounds n = 7 mice). j. Reaction times to hits and FA in task 2 (blue: cochlear implants n = 6 mice, red: cortical implants n = 7 mice, black: sounds n = 7 mice). k, l, m, n Proportion of licks to different rates of amplitude modulation at 45, 50, 55 and 60 dB SPL background noise respectively (blue: cochlear implants n = 6 mice, red: cortical implants n = 7 mice, black: sounds n = 7 mice). o, p Reaction times to hits and FA in task 4a and 4b (blue: cochlear implants n = 6 mice, red: cortical implants n = 7 mice, black: sounds n = 7 mice).

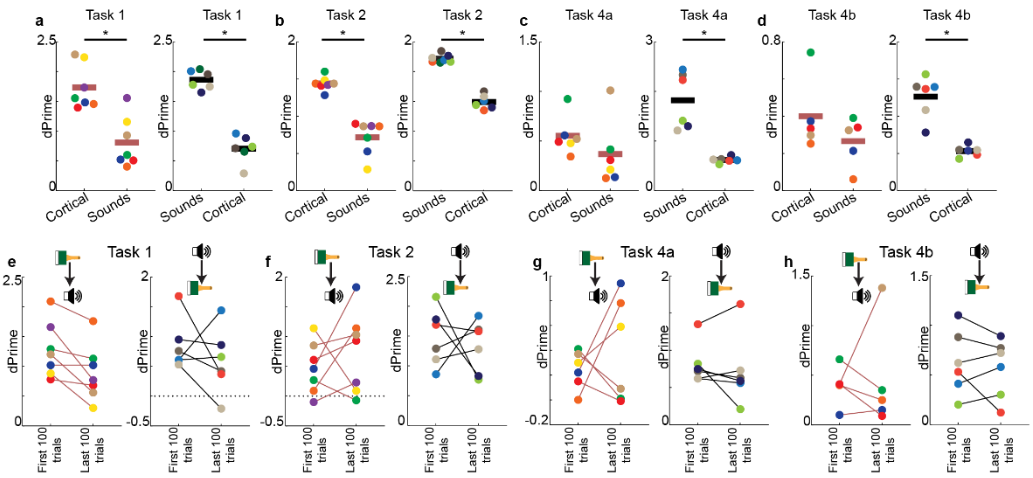


**Extended figure 4: Cross-modal performance is not explained by rapid relearning**

a-d Performance of mice (d') on last day of training to cortical implants (left) or sounds (right) and on testing day of opposite modality (red: cortical→sounds n = 7 for tasks 1 and n = 6 for task 4a and n = 5 for task 4b, black sounds → cortical n = 6, *, $p < 0.05$). e-h Performance of mice (d') on first 100 trials and last 100 trials of the cross-modal testing session (red: cortical→sounds n = 7 for tasks 1 and n = 6 for task 4a and n = 5 for task 4b, black sounds → cortical n = 6)

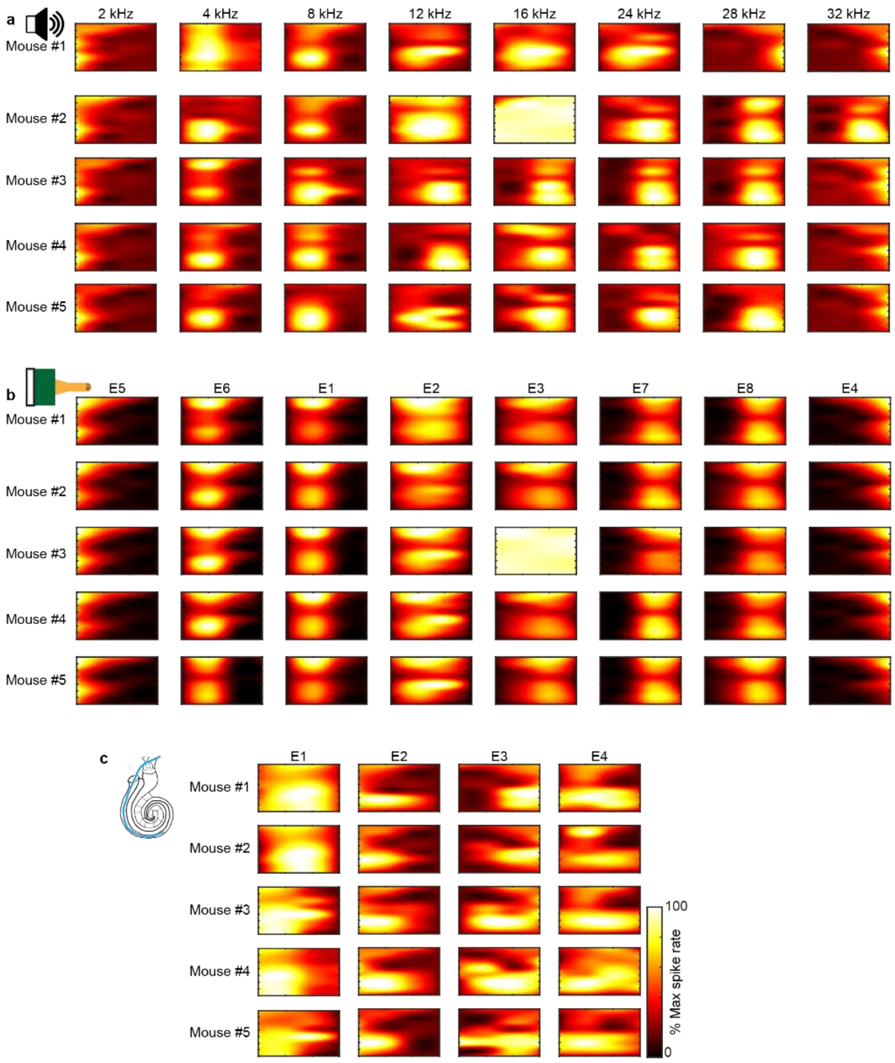


**Extended figure 5: Neuronal responses in A1.**

a. Example heatmaps of z-scored neuronal activity in A1 in response to individual pure tones for individual mice. b. Example heatmaps of z-scored neuronal activity in A1 in response to individual cortical electrode stimulations for individual mice. c. Example heatmaps of z-scored neuronal activity in A1 in response to cochlear individual electrode stimulations for individual mice.

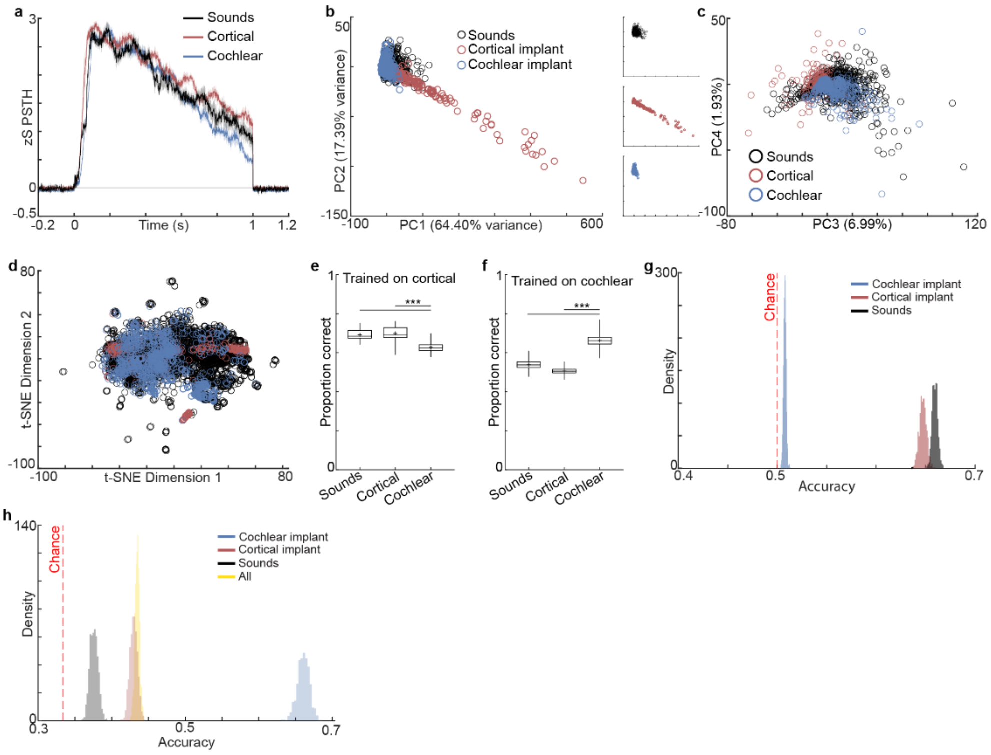


**Extended figure 6: Cortical stimulation evokes population responses more similar to natural sound than cochlear stimulation in task 1**

a. Average PSTH response to single electrodes/frequencies (black: sounds n = 203 cells from N = 5 mice, red: cortical n = 149 cells from N = 5 mice, blue: cochlear n = 129 cells from N = 5 mice). b. PCA of neuronal population responses to individual pure-tone frequencies onto components 1 and 2 (black, n = 203 cells from N = 5 mice), single-electrode cortical stimulation (red, n = 149 cells from N = 5 mice), and cochlear stimulation (blue, n = 129 cells from N = 5 mice). c. PCA of neuronal population responses to individual pure-tone frequencies onto components 3 and 4 (black, n = 203 cells from N = 5 mice), single-electrode cortical stimulation (red, n = 149 cells from N = 5 mice), and cochlear stimulation (blue, n = 129 cells from N = 5 mice). d. t-SNE visualization of trial-averaged neuronal responses for acoustic (black), cortical (red), and cochlear (blue) stimuli, based on the same dataset as in (b). e. Classification accuracy of a two-way SVM trained to distinguish spontaneous from cortical stimulation-evoked activity, and tested on held-out cortical stimulation responses, sound responses, and cochlear stimulation responses (***, $p < 0.001$). e. Classification accuracy of a two-way support vector machine (SVM) trained to distinguish spontaneous from cochlear stimulation-evoked activity, and tested on held-out cochlear stimulation responses, sound responses, and cortical stimulation responses (***, $p < 0.001$). g. Bootstrap distributions of two-way decoder accuracy. Each distribution shows the estimated decoding accuracy via bootstrap resampling (n = 1000). h. Bootstrap distributions of three-way decoder accuracy. Each distribution shows the estimated decoding accuracy via bootstrap resampling (n = 1000).

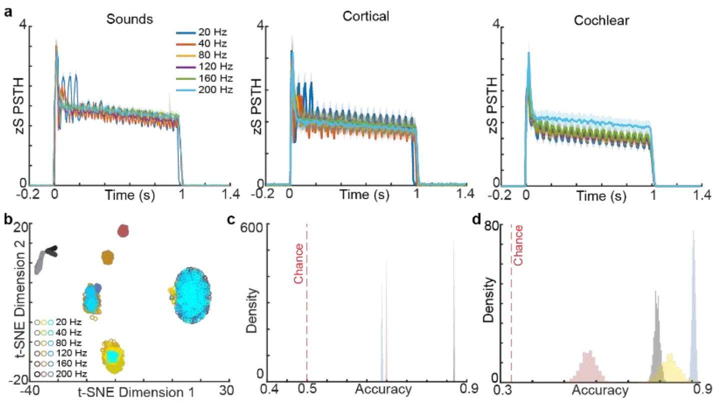


**Extended figure 7: Cortical stimulation preserves population-level coding of amplitude modulation.**

a. Average PSTH responses to different AM rates under sound (left), cortical stimulation (center), and cochlear stimulation (right) conditions. b. t-SNE visualization of trial-averaged neuronal responses for sound (black), cortical (red), and cochlear (blue) stimuli. c. Bootstrap distributions of two-way decoder accuracy. Each distribution shows the estimated decoding accuracy via bootstrap resampling (n = 1000). d. Bootstrap distributions of three-way decoder accuracy. Each distribution shows the estimated decoding accuracy via bootstrap resampling (n = 1000).

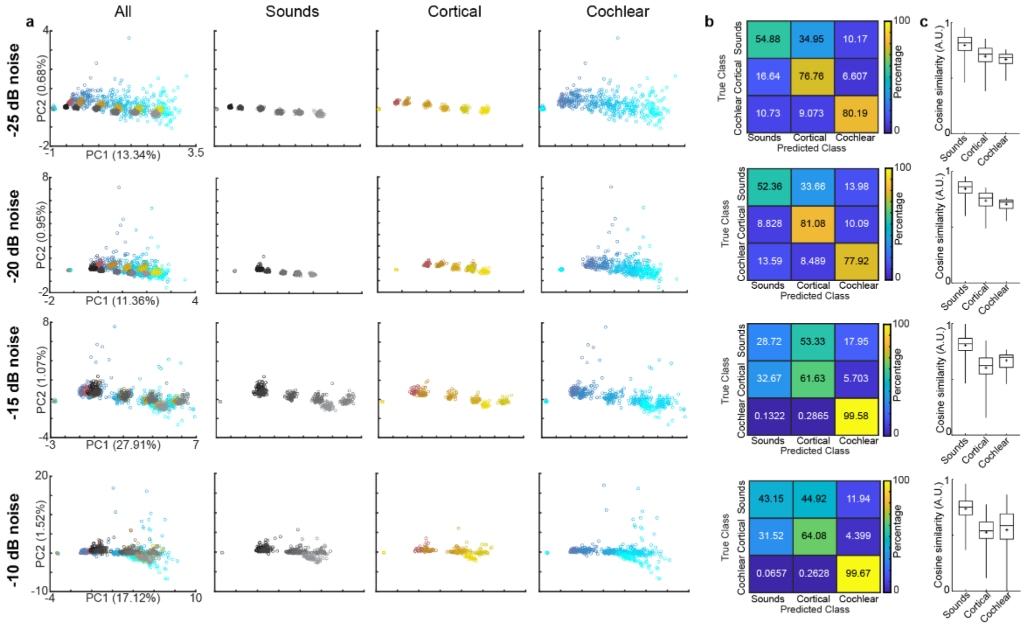


**Extended figure 8: Analysis of amplitude modulation in noise.**

a. PCA of responses AM stimuli delivered via sound (black), cortical stimulation (red), or cochlear stimulation (blue) with different levels of background noise. b. Confusion matrix from three-way SVM classification of AM responses in noise (training: n = 337 cells, testing n = 144 cells). c. Cosine similarity between trial-averaged auditory cortical population responses evoked by acoustic, cortical or cochlear AM stimulation in noise.

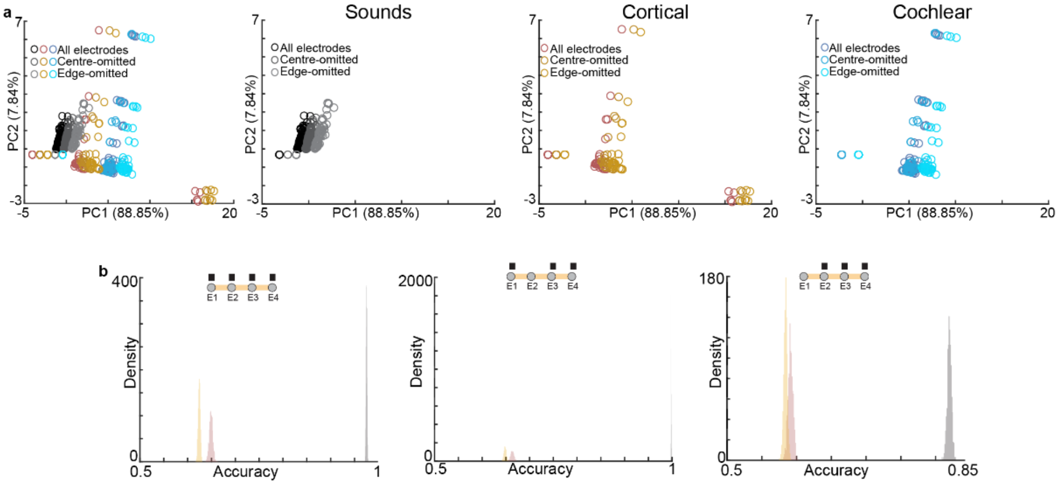


**Extended figure 9: Analysis of multi-frequency/electrode stimulation.**

a. PCA of responses multi-frequency/electrode stimuli delivered via sound (right, black), cortical stimulation (centre, red), or cochlear stimulation (right, blue). b. Bootstrap distributions of two-way decoder accuracy. Each distribution shows the estimated decoding accuracy via bootstrap resampling (n = 1000).
